# Top-down control of water intake by the endocannabinoid system

**DOI:** 10.1101/729970

**Authors:** Zhe Zhao, Edgar Soria-Gómez, Marjorie Varilh, Francisca Julio-Kalajzić, Astrid Cannich, Adriana Castiglione, Léonie Vanhoutte, Alexia Duveau, Philippe Zizzari, Anna Beyeler, Daniela Cota, Luigi Bellocchio, Arnau Busquets-Garcia, Giovanni Marsicano

**Author notes:** Correspondence should be addressed to: Giovanni Marsicano DVM, PhD, NeuroCentre Magendie, INSERM U1215 Université Bordeaux, Group Endocannabinoids and Neuroadaptation, 146 rue Léo-Saignat 33077 Bordeaux Cedex- France. Senior author (Luigi Bellocchio, Arnau Busquets-Garcia, Giovanni Marsicano).

## Abstract

Water intake is regulated by neocortical top-down circuits, but their identity and the cellular mechanisms involved are scantly known. Here, we show that endogenous activation of type-1 cannabinoid receptors (CB_1_) promotes water intake and that endocannabinoid modulation of excitatory projections from the anterior cingulate cortex to the basolateral amygdala is sufficient to guarantee physiological drinking. These data reveal a new circuit involved in the homeostatic control of water intake.

---

Water intake is crucial for maintaining body fluid homeostasis and animals’ survival^1^. Complex brain processes trigger thirst and drinking behavior, which arise upon losing blood volume (i.e. extracellular dehydration) or increasing blood osmolality (i.e. intracellular dehydration)^1^. However, the central mechanisms promoting water intake are still poorly understood. In the brain, the anterior wall of the third ventricle formed by the subfornical organ (SFO), the median preoptic nucleus, and the organum vasculosum of the lamina terminalis (OVLT) constitutes the primary structure sensing thirst signals and promoting water intake^2,3^. These subcortical regions are connected with the neocortex^1^. In particular, insular and anterior cingulate cortices (IC and ACC, respectively) have been shown to receive indirect projections from the SFO and OVLT in rats^4^, and water consumption after dehydration decreases ACC activity in humans^5^. Furthermore, recent evidence shows that stimulation of the anterior part of IC promotes drinking behavior, whereas stimulation of the posterior part exerts the opposite effect^6^. These studies highlight the importance of cortical regions in the regulation of water intake^1,4-6^.

Type-1 cannabinoid receptors (CB_1_) are widely and abundantly expressed in the central nervous system where they modulate a variety of functions, including feeding behavior^7-9^. However, the role of CB_1_ receptors in the control of water intake is still a matter of debate, since pharmacological activation or blockade of CB_1_ receptors produced contradictory results in drinking behavior experiments^10,11^. In this study, we identified a novel and specific cortical circuit where CB_1_ receptors modulate water intake.

To examine the role of CB_1_ receptors in the control of water intake, we first tested *CB*_*1*_ knockout mice (*CB*_*1*_-KO)^12^ under different experimental conditions. No significant difference was observed between *CB*_*1*_ wild-type (*CB*_*1*_*-*WT) and *CB*_*1*_-KO littermates in daily water intake (**Supplementary Fig. 1a**). However, *CB*_*1*_-KO mice drunk less than WT littermates after 24-hour water deprivation (**Fig. 1a, Supplementary Fig. 1b**), without any change in food intake (**Supplementary Fig. 1c**), indicating that CB_1_ receptors participate in water deprivation-induced drinking behavior. Water deprivation triggers both intracellular and extracellular dehydration that can lead to water intake through different pathways^1^. To discriminate the impact of CB_1_ receptor signaling on either of these mechanisms, we first applied systemic (intraperitoneal, IP) or intracerebroventricular (ICV) injections of sodium chloride (NaCl), which are known to induce water intake by mimicking intracellular dehydration^1^. As compared to wild-type littermates, mice lacking CB_1_ receptors displayed a lower water intake induced by both IP or ICV NaCl administration (**Fig. 1b, c, Supplementary Fig. 1d**). Extracellular dehydration promotes the production of angiotensin II (ANG), which can induce drinking behavior and salt appetite^1^. Thus, to mimic this condition, mice received ICV injections of ANG. Notably, the ANG-induced water intake was blunted in *CB*_*1*_-KO mice **(Fig. 1d)**, indicating that endocannabinoid signaling controls drinking behavior induced by both intracellular and extracellular dehydration mechanisms. Importantly, the acute systemic pharmacological blockade of CB_1_ receptors decreased drinking under water deprivation and NaCl injections (**Fig. 1e, f**), indicating that endocannabinoid signaling is required at the moment of drinking and that the phenotype of *CB*_*1*_-KO mice is not due to the long-lasting deletion of the gene^13^. Concomitantly with the abundant brain expression, CB_1_ receptors are also present in peripheral organs^7^, suggesting that peripheral control of body water levels or blood osmolality might underlie the endocannabinoid-dependent regulation of water intake. However, measurements of body water composition and blood osmolality did not reveal any difference between *CB*_*1*_-KO mice and *CB*_*1*_-WT littermates (**Supplementary Fig. 1e, f**). Altogether, these results indicate that endogenous activation of CB_1_ receptors contributes to drinking behavior induced by both intracellular and extracellular dehydration conditions, likely through central mechanisms.

**Figure 1.**
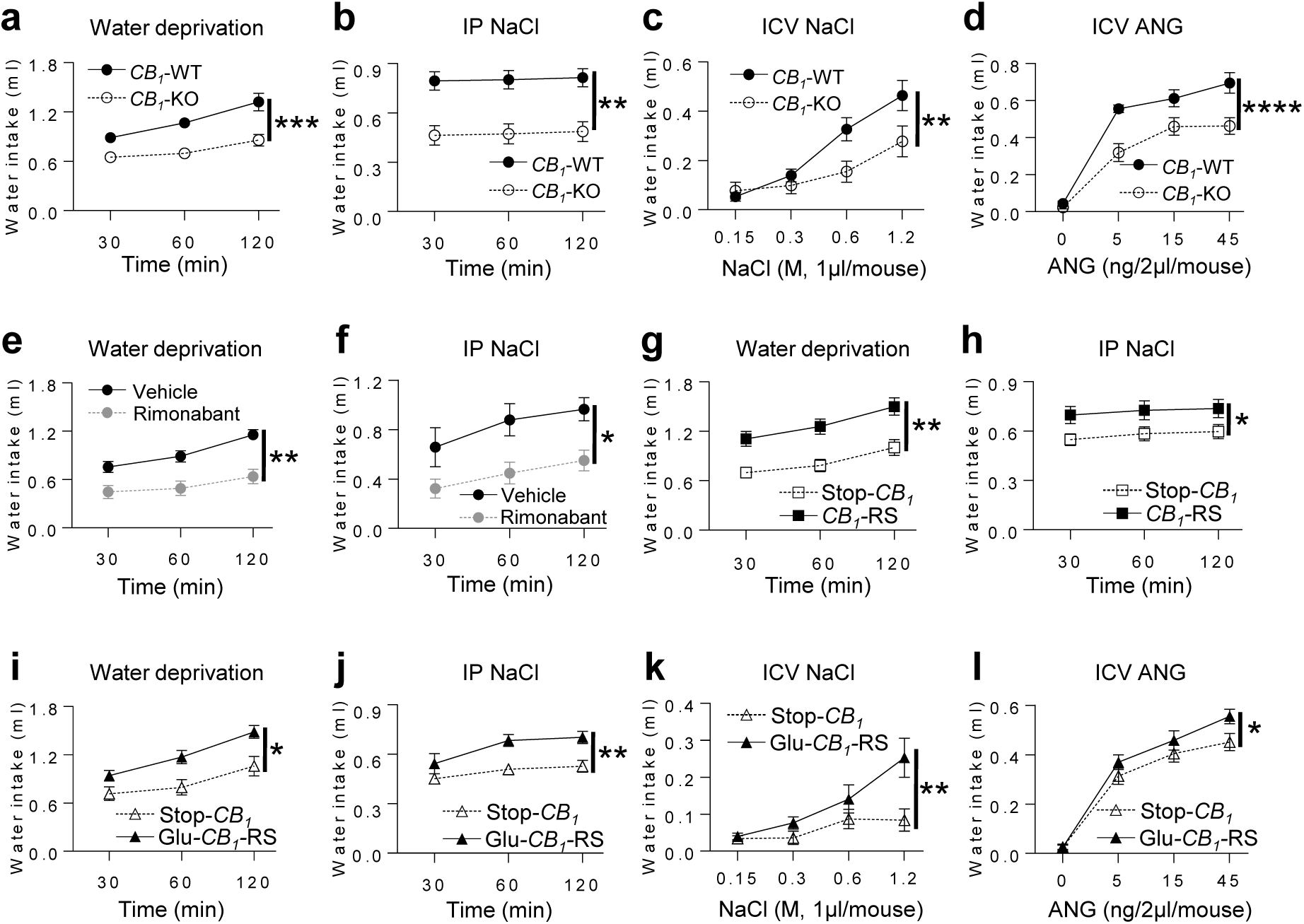
Global deletion of CB_1_ decreases water intake induced by different dehydrations, whereas re-expression of *CB*_*1*_ in cortical glutamatergic neurons is sufficient to promote water intake. **a-d**, Cumulative water intake of *CB*_*1*_-WT (Black circles) and *CB*_*1*_-KO (Open circles) mice after 24-hour water deprivation (*CB*_*1*_-WT n=10, *CB*_*1*_-KO n=8), IP 1M NaCl, 10ml/kg body weight (*CB*_*1*_-WT n=10, *CB*_*1*_-KO n=8), ICV NaCl (*CB*_*1*_-WT n=13, *CB*_*1*_-KO n=10), and ICV ANG (*CB*_*1*_-WT n=11, *CB*_*1*_-KO n=13). **e-f**, Cumulative water intake induced by 24-hour water deprivation (Vehicle n=9, Rimonabant n=10) or IP 1.5M NaCl, 10ml/kg body weight (Vehicle n=6, Rimonabant n=7) after systemic blockade of CB_1_ receptors (Rimonabant, 3mg/kg, gray circles; Vehicle, black circles). **g-h**, Cumulative water intake induced by 24-hour water deprivation (Stop-*CB*_*1*_ n=9, *CB*_*1*_-RS n=12), IP 1M NaCl, 10ml/kg body weight (Stop-*CB*_*1*_ n=9, *CB*_*1*_-RS n=11) in Stop-*CB*_*1*_ (Open squares) and *CB*_*1*_-RS (Black squares) mice. **i-l**, Cumulative water intake induced by 24-hour water deprivation (Stop-*CB*_*1*_ n=11, Glu-*CB*_*1*_-RS n=11), IP 1M NaCl, 10ml/kg body weight (Stop-*CB*_*1*_ n=11, Glu-*CB*_*1*_-RS n=11), ICV NaCl (Stop-*CB*_*1*_ n=13, Glu-*CB*_*1*_-RS n=11), and ICV ANG (Stop-*CB*_*1*_ n=15, Glu-*CB*_*1*_-RS n=13) in Stop-*CB*_*1*_ (Open triangles) and Glu-*CB*_*1*_-RS (Black triangles) mice. All data are showed as mean ± s.e.m, and were statistically analyzed by the two-way repeated measures ANOVA, \**P* < 0.05, \*\**P* < 0.01, \*\*\**P* < 0.001, \*\*\*\**P* < 0.0001.

CB_1_ receptors are expressed in many different brain regions and in distinct cell types^7,8,13^. To identify the specific cell-type involved in CB_1_ receptor-dependent control of water intake, we used conditional mutant mice carrying deletion of the *CB*_*1*_ gene in specific cell types, such as cortical glutamatergic neurons (Glu-*CB*_*1*_-KO)^14,15^, forebrain GABAergic neurons (GABA-*CB*_*1*_-KO)^14,15^, glial fibrillary acidic protein-positive cells (mainly astrocytes, GFAP-*CB*_*1*_-KO)^14,16^ and dopamine receptor D_1_-positive cells (D_1_-*CB*_*1*_-KO)^14,17^. All these cell types have been implicated in the control of water intake^1-3^. Surprisingly, however, none of these mutant lines displayed significant phenotypes in drinking behavior induced by water deprivation or NaCl injections (**Supplementary Fig. 2a-h**).

It is particularly puzzling how global, but not cell type-specific, CB_1_ deletion can impact water intake. This may be due to the redundancy of CB_1_ receptor-dependent pathways controlling a function as vital as water intake. In this context, despite the general necessary role of the endocannabinoid system in controlling drinking behavior, this redundancy would decrease the specific *necessity* of selected subpopulations of CB_1_ receptors. This, however, does not exclude that CB_1_ receptor-dependent control of specific cell populations might play *sufficient* roles in controlling stimulated water intake. To address this possibility, we adopted a rescue approach and we used mice carrying specific and exclusive re-expression of the CB_1_ protein in specific cell types (Stop-*CB*_*1*_ mice approach)^18,19^. A “floxed-stop” cassette prevents the expression of CB_1_ receptors in the stop-*CB*_*1*_ mutant line, similarly as in global *CB*_*1*_-KO mice. Viral or transgenic expression of the Cre recombinase, however, induces the re-expression of the CB_1_ receptors in particular brain regions and/or cell types over a “knockout-like” background^18,19^.

First, we verified that Stop-*CB*_*1*_ mice displayed the same impaired water intake as *CB*_*1*_-KO mice and that global re-expression of the CB_1_ protein is able to fully rescue water intake under deprivation and NaCl injections (*CB*_*1*_-RS for CB_1_ rescued ; **Fig. 1g,h**)^18,19^. Re-expression of CB_1_ protein in GABAergic neurons (GABA-*CB*_*1*_-RS mice)^18^, which include the large majority of brain CB_1_ receptors^7,8,13^, did not rescue drinking behavior either after water deprivation or IP NaCl injection (**Supplementary Fig. 2i, j**). Interestingly, however, re-expression of CB_1_ receptors in cortical glutamatergic neurons (Glu-*CB*_*1*_-RS)^19^, which represents a minority of the receptor in the brain^7,8,13^, significantly rescued water intake induced by water deprivation, by systemic or central injection of NaCl, or by ICV ANG administration (**Fig. 1i-l**). These data indicate that the presence of CB_1_ receptors in cortical glutamatergic neurons is sufficient to promote water intake induced by different conditions.

Amongst other neocortical areas, the insular cortex (IC) has been directly shown to regulate water intake^6^. Therefore, we tested whether specific re-expression of CB_1_ receptors in this brain region might rescue the impairment of water intake observed in Stop-*CB*_*1*_ mice. Multiple local injections of an adeno-associated virus expressing Cre recombinase (AAV-Cre) into the IC of Stop-*CB*_*1*_ mice resulted in a consistent *CB*_*1*_ re-expression in both anterior and posterior portions of this brain region (IC-*CB*_*1*_-RS; **Supplementary Fig. 3a-e**). However, this manipulation did not rescue the water intake associated with lack of CB_1_ receptor protein (**Supplementary Fig. 3f,g**). Recent evidence points to the idea that the anterior and posterior parts of the IC play opposite roles in the control of drinking behavior^6^. In particular, activation of neurons located in the anterior IC (aIC) increases water intake, whereas the same manipulation of the posterior IC (pIC) exerts the opposite effect^7^. Considering that activation of CB_1_ receptors generally reduces neuronal activity^10^, we reasoned that endocannabinoid control of the pIC leads to decreased neuronal activity and promotes drinking behavior. To test this possibility, we re-expressed the CB_1_ protein exclusively in the pIC of Stop-*CB*_*1*_ mice (pIC-*CB*_*1*_-RS, **Supplementary Fig. 4a,b and 3e**), where the lack of the receptor should logically induce a reduction of drinking. However, also this partial re-expression did not rescue the phenotype of Stop-*CB*_*1*_ mice (**Supplementary Fig. 4c,d**), strongly suggesting that CB_1_ receptors in this brain region do not play a major role in water intake.

Recent studies suggest that the anterior cingulate cortex (ACC) might participate in the regulation of water intake^1,4,5^. Using a similar approach as above, we generated ACC-*CB*_*1*_-RS mice, in which the CB_1_ protein is re-expressed only in ACC principal neurons (**Fig. 2a-c**). Notably, ACC-*CB*_*1*_-RS mice displayed significantly higher water intake than Stop-*CB*_*1*_ littermates (ACC-*CB*_*1*_-SS) both after water deprivation and IP NaCl injection (**Fig. 2d,e**), indicating that the presence of CB_1_ receptors in principal neurons of the ACC is sufficient to promote drinking behavior induced by water deprivation and NaCl treatment.

**Figure 2.**
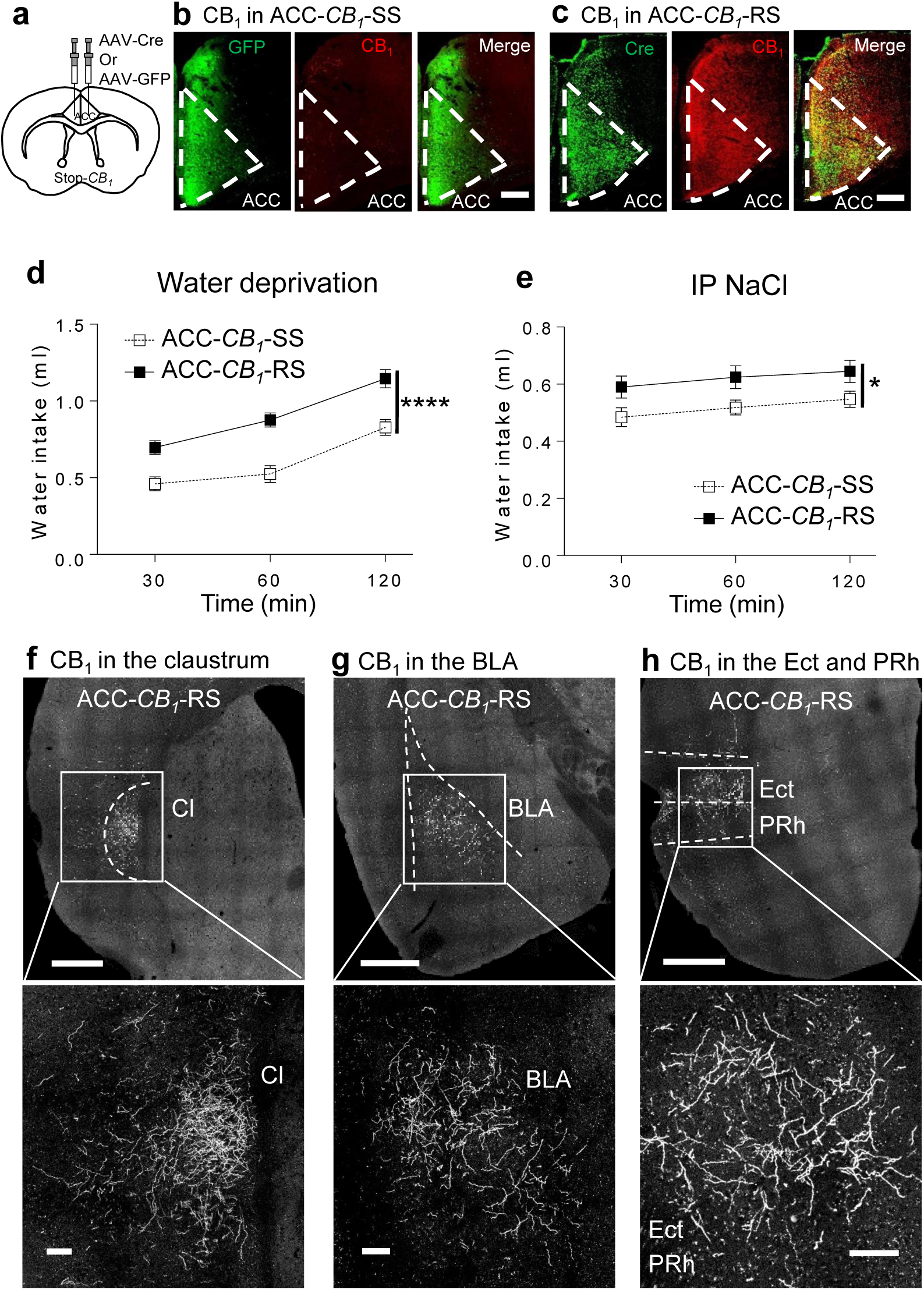
Re-expression of CB_1_ in the ACC is sufficient to promote water intake. **a**, Schematic representation of CB_1_ rescue approach in the ACC of Stop-*CB*_*1*_ mice. **b-c**, CB_1_ (red) immunostaining in the ACC of ACC-*CB*_*1*_-SS and ACC-*CB*_*1*_-RS, respectively. Scale bar, 200 μm. **d-e**, Cumulative water intake of ACC-*CB*_*1*_-SS (Open squares) and ACC-*CB*_*1*_-RS (Black squares) mice after 24-hour water deprivation (ACC-*CB*_*1*_-SS n=17, ACC-*CB*_*1*_-RS n=20) or IP 1M NaCl, 10ml/kg body weight (ACC-*CB*_*1*_-SS n=18, ACC-*CB*_*1*_-RS n=20). **f-h**, Presynaptic CB_1_ receptors located in the CI, BLA and Ect/PRh in a ACC-*CB*_*1*_-RS mouse. Scale bar, 500 μm and 100 (Amplified images) μm. All data are showed as mean ± s.e.m, and were statistically analyzed by the two-way repeated measures ANOVA, *\*P* < 0.05, *\*\*\*\*P* < 0.0001.

As the ACC is a heterogeneous structure targeting multiple downstream regions, we next aimed at identifying which CB_1_-positive projections from ACC are responsible for the stimulation of drinking behavior. First, we mapped the ACC neuronal projections interested by our local viral treatments. The injection of an AAV-CaMKIIα-GFP virus into the ACC revealed that principal neurons of this neocortical region project to many brain areas, including the basolateral amygdala (BLA), the claustrum (Cl), the medial caudate putamen, the lateral habenula (**Supplementary Fig. 5 and video 1**). In order to analyze the expression of presynaptic CB_1_ receptors in these ACC projections, we evaluated the distribution of the CB_1_ protein in ACC-*CB*_*1*_-RS mice. Interestingly, CB_1_ receptors were mainly present in the claustrum, the BLA, as well as in the ectorhinal and perirhinal cortices (**Fig. 2f-h, Supplementary video 2**). Recent evidence actually indicates that BLA is involved in the control of drinking behavior^6,20^. We therefore asked whether CB_1_ receptors expressed in ACC to BLA projections (ACC-BLA) are sufficient to promote water intake (**Fig. 3a**). To obtain selective rescue of the CB_1_ protein in ACC-BLA terminals, we used a retrograde viral approach in the Stop-*CB*_*1*_ mice. The injection of a retrograde AAV (rAAV2-retro) expressing flippases coupled to blue fluorescent protein (rAAV2-retro-FLIPo-EBFP) into the BLA of Stop-*CB*_*1*_ mice was associated with the simultaneous infusion of another AAV carrying a FLIPo-dependent expression of Cre recombinase (AAV-FRT-iCre) into the ACC (**Fig. 3b**). These combinatorial viral manipulations resulted in a strong re-expression of CB_1_ protein in ACC-BLA projecting neurons of Stop-*CB*_*1*_ mice (ACC-BLA-*CB*_*1*_-RS mice; **Fig. 3c-e**). Strikingly, after water deprivation and IP NaCl injection, ACC-BLA-*CB*_*1*_-RS mice consumed significantly more water than control mice (**Fig. 3f,g**), revealing the key role of CB_1_ receptor-dependent control of drinking in this specific brain circuit.

**Figure 3.**
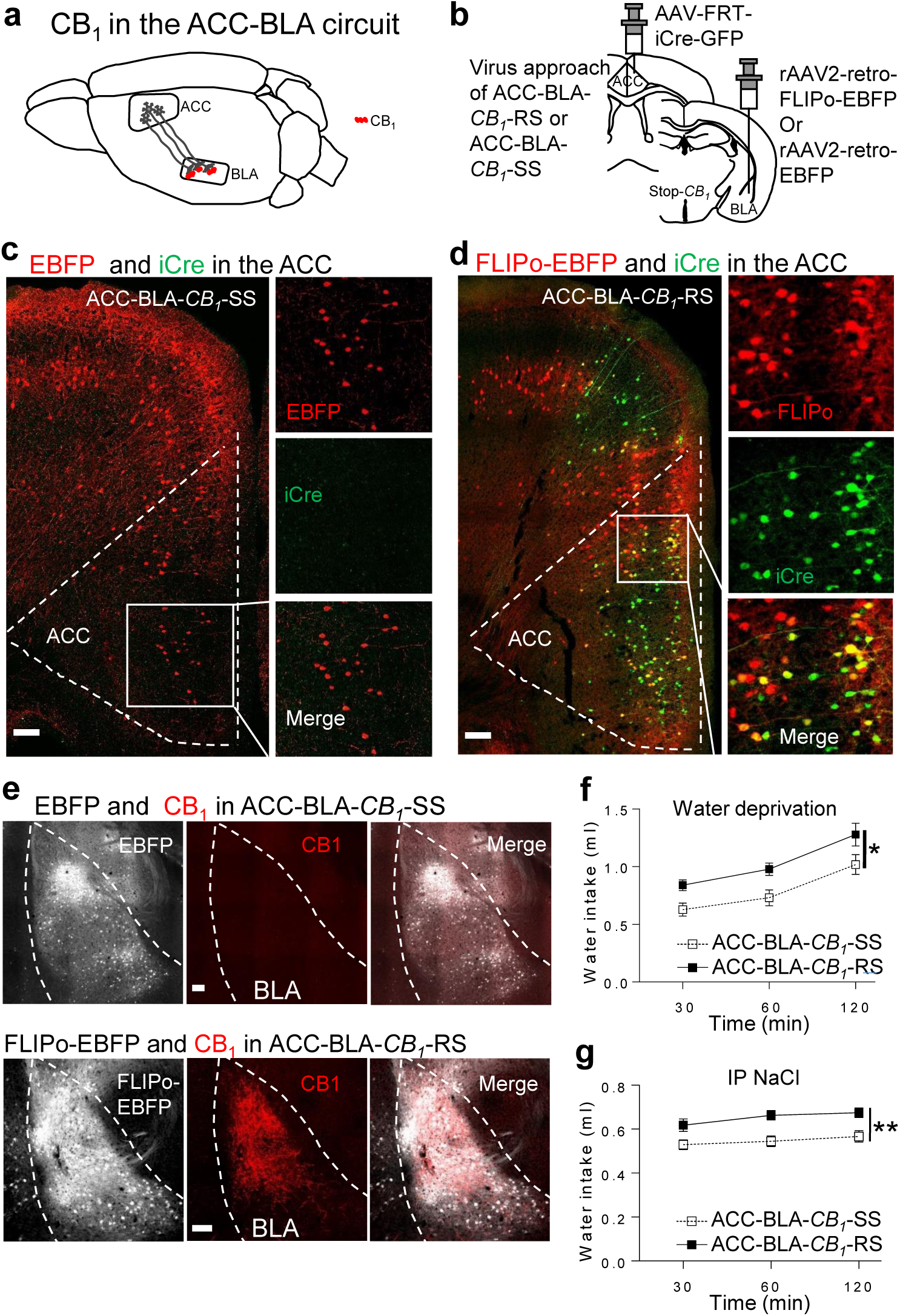
CB_1_ receptors located in ACC-BLA is sufficient to promote water intake. **a**, Schematic representation of CB_1_ receptors located in the ACC-BLA circuit. **b**, Viral approach to specifically rescue *CB*_*1*_ in the ACC-BLA circuit. **c**, EBFP (pseudo red) and iCre-GFP (green) in ACC sections of ACC-BLA-*CB*_*1*_-SS. Scale bar, 100 μm. **d**, FLIPo-EBFP (pseudo red) and iCre-GFP (green) in ACC section of ACC-BLA-*CB*_*1*_-RS. Scale bar, 100 μm. **e-f**, CB_1_ (red) immunostaining in BLA section of ACC-BLA-*CB*_*1*_-SS and ACC-BLA-*CB*_*1*_-RS. Scale bar, 100 μm. **g-h**, Cumulative water intake of ACC-BLA-*CB*_*1*_-SS (Open squares) and ACC-BLA-*CB*_*1*_-RS (Black squares) mice after 24-hour water deprivation (ACC-BLA-*CB*_*1*_-SS n=10) and 1M IP NaCl, 10ml/kg body weight (ACC-BLA-*CB*_*1*_-SS n=12). All data are showed as mean ± s.e.m, and were statistically analyzed by the two-way repeated measures ANOVA, *\*P* < 0.05, *\*\*P* < 0.01.

This study reveals an unforeseen circuit mechanism for top-down control of a fundamental life function such as water intake. Specifically, general CB_1_ receptor activity is necessary to promote water intake, and its control of ACC principal neurons impinging onto BLA is sufficient to promote drinking behavior. These data highlight the complexity of brain control of water intake and underline the importance of top-down regulatory circuits in these processes.

## Author contributions

Z.Z. and G.M. conceived the project. Z.Z. and G.M. designed the experiments and analyzed data with the input of E.S., L.B., and A.B. Z.Z. performed the experiments and collected data. Z.Z., L.B., and G.M. wrote the manuscript. M.V. and F.J. performed immunohistochemistry experiments. A.C., A.C., L.V., A.D., and P.Z. assisted in performing experiments. A.B. and D.C. discussed the study. All authors read and edited the manuscript.

## Acknowledgments

We thank the animal facility and the genotyping platform of the NeuroCentre Magendie (INSERM U1215 Unit) for providing assistance in the animal breeding, maintenance and genotyping. We also thank Drs. Aude Panatier and Stéphane Oliet of NeuroCentre Magendie for providing the Osmometer. We thank the Bordeaux Imaging Center, a service unit of the CNRS-INSERM and Bordeaux University, member of the national infrastructure France BioImaging supported by the French National Research Agency (ANR-10-INBS-04), for providing the confocal microscope (Leica TCS SP8), the slide scanner (Nanozoomer 2.0HT, Hamamatsu Photonics France), and Imaris software (Imaris, Oxford instrument, UK), the help of Sébastien Marais is acknowledged. HHMI Janelie farm research campus is acknowledged for providing the rAAV2-retro helper. We thank the Viral Vector Facility (VVF) of Neuroscience Center Zurich (ZNZ) for providing the rAAV2-retro-FLIPo-EBFP and the rAAV2-retro-EBFP viral vectors. We also thank Dr. Karl Deisseroth from Stanford University, Stanford, CA for providing the plasmid of AAV-CaMKIIα-GFP. This work is supported by the China Scholarship Council (to Z.Z.), INSERM (to G.M., D.C., A.B.), Nouvelle Aquitaine Region (to D.C., G.M.), European Research Council (Endofood, ERC–2010–StG–260515 and CannaPreg, ERC-2014-PoC-640923, MiCaBra, ERC-2017-AdG-786467, to G.M.), Fondation pour la Recherche Medicale (FRM, DRM20101220445, to G.M.), the Human Frontiers Science Program, Region Aquitaine, Agence Nationale de la Recherche (ANR, NeuroNutriSens ANR-13-BSV4-0006, ORUPS ANR-16-CE37-0010-01 and CaCoVi ANR-18-CE16-0001-02, to G.M.) and BRAIN ANR-10-LABX-0043, to G.M.

## Supplementary information

### Supplementary online methods

#### Mice

All experiments were approved by the Committee on Animal Health and Care of INSERM and the French Ministry of Agriculture and Forestry. The authorizing number from the ethical committee is 15493. Maximal efforts were made to reduce the suffering and the number of used mice. All behavioral experiments were performed during the light phase and animals were kept in individual cages under standard conditions in a day/night cycle of 12/12 hours (lights on at 7 am). Male wild-type C57BL/6n mice purchased from Janvier (France) were used for the pharmacological experiments. All the used mutant mice were generated and identified in previous studies, e.g. global CB_1_ knockout (*CB*_*1*_-KO) mice^12^, deletion of CB_1_ receptors is specific in cortical glutamatergic Nex positive neurons (Glu-*CB*_*1*_-KO)^14,15^, forebrain GABAergic Dlx5/6 positive neurons (GABA-*CB*_*1*_-KO)^14,15^, astrocytes (GFAP-*CB*_*1*_-KO)^14,16^, and dopamine receptor type 1 positive neurons (D_1_-*CB*_*1*_-KO) ^14,17^. The stop-*CB*_*1*_ mice^18,19^ (lack of CB_1_), global re-expression of CB_1_ receptors (*CB*_*1*_-RS)^18,19^, re-expression of CB_1_ receptors is specific in forebrain GABAergic Dlx5/6 positive neurons (GABA-*CB*_*1*_-RS)^18^ and cortical glutamatergic Nex positive neurons (Glu-*CB*_*1*_-RS)^19^. The mice used in this study were 7-10 weeks old at the beginning of the experiments.

#### Water intake assays

Water intake was observed at 30, 60 and 120 minutes after 24-hour water deprivation and intraperitoneal (IP) injection of 1M sodium chloride (NaCl, VWRV0241) with 10ml/kg body weight. In the pharmacological experiments, Rimonabant (3mg/kg, 9000484, Cayman Chemical Company US) and vehicle (4% ethanol, 4% Cremophor, 92% saline) were injected half an hour prior to the water intake test of the water deprivation or IP injection of 1.5 M NaCl with 10ml/kg body weight. For the mice of ICV injection, water intake was observed at 30 minutes after intracerebroventricular (ICV) injection of Angiotensin **II** (ANG, Bachem, H-1705.0025) and NaCl. We started to test water intake 7 days after the ICV cannula implantation. ICV injection was once a day in each mouse. In the progressive ANG dose-response experiments, we did ICV injections of saline, 5 ng, 15 ng, and 45 ng ANG (2μl/mouse) in different days. Then, we start the ICV NaCl injection 3 days after the last ICV ANG injection, ICV injection was once a day in each mouse. In the progressive NaCl dose-response experiments, we did ICV NaCl injections of 0.15M, 0.3M, 0.6M, and 1.2M NaCl (1μl/mouse) in the different days. In order to make sure that mice were drinking normally before the treatments, each mouse was observed the daily water intake during the experiments.

#### Body water composition analysis

The basal body water composition test was performed in mice by using a mouse-specific nuclear magnetic resonance whole body composition analyzer (EchoMRITM-900, EchoMedical Systems, Houston, TX). Mice were placed in a specific chamber without strong movements, each readout was done within 1 minute. Mice were put back to home cages after the test.

#### Plasma osmolality analysis

Plasma osmolality was tested by Osmometer 3320 (Advanced Instruments, France). Facial veil blood collection was applied in this experiment. Blood was collected and put in the Micro tube 1.3 ml K3E (SARSTEDT, 41.1395.005), then blood samples were remained in room temperature for 30 minutes. By using a refrigerated centrifuge (VWR Micro Star 17R), blood samples were centrifuged with 4000 rpm for 15 minutes at 4°C. Following centrifugation, the plasma was immediately transferred to a clean eppendorf tube and put on the ice for the osmolality test.

#### Surgery and viral administration

Mice were anesthetized by isoflurane (5% induction, then, 2% during the surgery) and placed on a stereotaxic apparatus (Model 900, KOPF instruments, CA, USA) with a mouse adaptor and lateral ear bars. For viral vectors delivery, AAV vectors were loaded in a glass pipette and fused by a pump (UMP3-1, World Precision Instruments, FL, USA). AAV-GFP (Hybrid AAV1/2, 5 x 10E10 vg/ml), AAV-Cre-GFP (Hybrid AAV1/2, 4.5 x 10E10 vg/ml) were injected into the insula (IC) (200nl/side, 100nl/min). The coordinate of anterior IC injection is AP +1.2mm, ML ± 3.0mm, DV 3.5mm, and the coordinate of posterior IC injection is AP −0.3mm, ML ± 3.7mm, DV 4.0mm. AAV-CaMKIIα-GFP (Hybrid AAV1/2, >1 x 10E10 vg/ml) or AAV-CaMKIIα-Cre-HA (Hybrid AAV1/2, >1 x 10E10 vg/ml. The plasmids were provided by Karl Deisseroth, Stanford University, Stanford, CA) were injected into the anterior cingulate cortex (ACC) (200nl/side, 100nl/min). The coordinate of ACC injection is AP +0.6mm, ML ± 0.3mm, DV 2.0mm, For the ACC-BLA-*CB*_*1*_-RS or ACC-BLA-*CB*_*1*_-SS mice, the AAV-FRT-iCre-GFP (Addgene #24593, ZNZ VVF v245, 6.3 x 10E12 vg/ml) was injected into ACC with the coordinates mentioned above in both group mice (200nl/side, 100nl/min). The rAAV2-retro-FLIPo-EBFP (Addgene #60663, ZNZ VVF v151, 6.4 x 10E12 vg/ml) or rAAV2-retro-EBFP (ZNZ VVF v140, 4.1 x 10E12 vg/ml) were injected into the BLA with the coordinates AP −1.6mm, ML ±3.3mm, DV 4.9 mm (150nl/side, 100nl/min). AAV-FRT-iCre-GFP, rAAV2-retro-FLIPo-EBFP, and rAAV2-retro-EBFP were produced by the Viral Vector Facility (VVF) of the Neuroscience Center Zurich (ZNZ). The re-expression of CB_1_ receptors was verified by the immunohistochemistry in all the mice used in the behavioral experiments. Above coordinates were according to the mouse brain in stereotaxic coordinates by Paxinos and Franklin, 2001 (Second edition).

#### Immunohistochemistry and imaging

After the behavioral experiment, mice were anesthetized with pentobarbital (Exagon, 400 mg/kg body weight), transcardially perfused first with the phosphate-buffered solution (PBS, 0.1M, pH 7.4), then fixed by 4% formaldehyde. After brain extraction, serial brain coronal sections were cut at 40 μm and collected in PBS at room temperature (RT). Sections were permeabilized in a blocking solution of 4% donkey serum, 0.3% Triton X-100 and 0.02% sodium azide prepared in PBS for 1 hour at RT. For the CB_1_ immunohistochemistry, free-floating sections were incubated with goat CB_1_ receptors polyclonal primary antibodies (CB_1_-Go-Af450-1; 1:2000, Frontier Science Co. ShinKO-nishi, Ishikari, Hokkaido, Japan) for 48 hours at 4°C. The antibody was prepared in the blocking solution. After three washes, the sections were incubated with a secondary antibody anti-goat Alexa Fluor 555 (A21432, 1:500, Fisher Scientific) for 2 hours at RT and then washed in PBS at RT. For the HA immunohistochemistry, it is similar with the CB_1_. Sections were incubated in anti-HA tag monoclonal antibody (1:1000, Fisher Scientific, 2-2.2.14) for 18 hours at 4°C and in secondary antibody anti-mouse Alexa Fluor 488 (A21202, 1:500, Fisher Scientific) for 2 hours at RT. All sections were mounted, dried and cover slipped. The sections were analyzed with a Nanozoomer microscope (Hamamatsu, Japan) and Leica SP8 confocal microscope (Leica, Germany). Images were analyzed by Image J (NIH). For the mouse brain reconstruction, images were collected by Nanazoomer, Z-stack images were made by Image J, and the 3D reconstruction and videos were made by Imaris software (Imaris, Oxford instrument, UK).

#### Statistics

Data handling and statistical analysis were performed using Microsoft Excel and GraphPad Prism 6 software. For the dose-response experiments of ICV NaCl and ICV ANG, and body water composition data were statistically analyzed by two-way analysis of variance (ANOVA). For the water intake test with several time points, data were statistically analyzed by the two-way repeated measures ANOVA. The data of IP NaCl dose response and plasma osmolality were statistically analyzed by two-tailed Student’s t-test. P values of ≤0.05 were considered statistically significant at a confidence interval of 95%. For detailed statistical analysis, see statistical tables (Supplementary tables 1-2).

**Supplementary figure 1.**
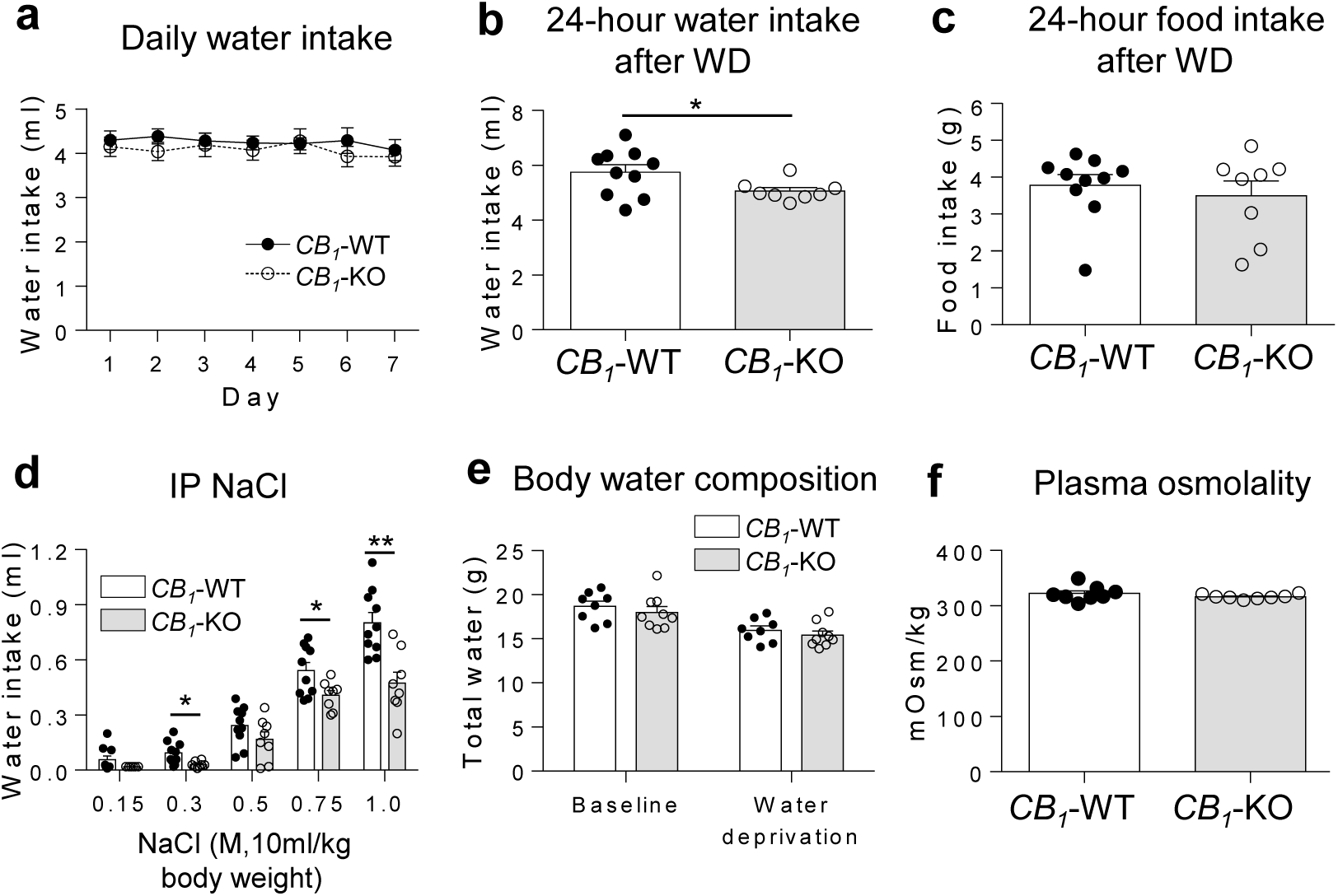
Decrease of stimulated water intake in *CB*_*1*_-KO mice is independent of food intake, body water composition, and plasma osmolality. **a**, Daily water intake of *CB*_*1*_-WT (Black circles, n=9) and *CB*_*1*_-KO (Open circles, n=8). **b**, Water intake in 24 hours after 24-hour water deprivation in *CB*_*1*_-WT (White, n=10) and *CB*_*1*_-KO (Gray, n=8) mice. **c**, Food intake in 24 hours after 24-hour water deprivation in *CB*_*1*_-WT (White, n=10) and *CB*_*1*_-KO (Gray, n=8) mice. **d**, Water intake in 1 hour after IP NaCl, 10ml/kg body weight at different doses in *CB*_*1*_-WT (White, n=10) and *CB*_*1*_-KO (Gray, n=8) mice. **e**, Body water composition test in *CB*_*1*_-WT (White, n=8) and *CB*_*1*_-KO (Gray, n=9) mice. **f**, Blood plasma osmolality test in *CB*_*1*_-WT (White, n=8) and *CB*_*1*_-KO (Gray, n=8) mice. All data are showed as mean ± s.e.m. Data of IP NaCl dose response and 24-hour water intake after water deprivation were statistically analyzed by two-tailed Student’s t-test. *\*P* < 0.05, *\*\*P* < 0.01.

**Supplementary figure 2.**
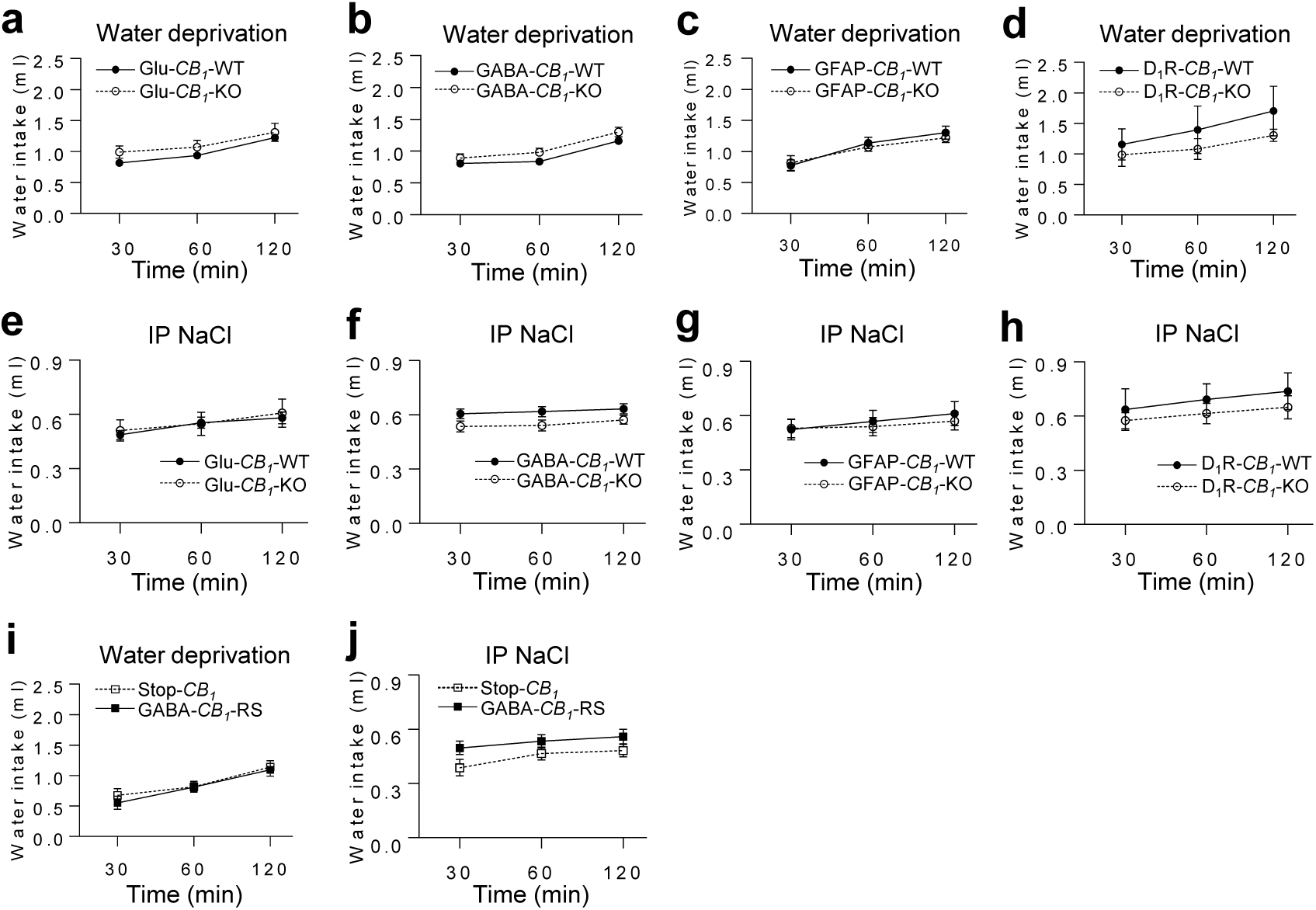
Deletion or re-expression of CB_1_ receptors in specific cell types did not affect water intake. **a-d**, Cumulative water intake after 24-hour water deprivation in Glu-*CB*_*1*_-WT (Black circles, n=18) and Glu-*CB*_*1*_-KO (Open circles, n=11), GABA-*CB*_*1*_-WT (Black circles, n=6) and GABA-*CB*_*1*_-KO (Open circles, n=10), GFAP-*CB*_*1*_-WT (Black circles, n=7) and GFAP-*CB*_*1*_-KO(Open circles, n=11), D_1_-*CB*_*1*_-WT (Black circles, n=5) and D_1_-*CB*_*1*_-KO(Open circles, n=7). **e-h**, Cumulative water intake after IP 1M NaCl, 10ml/kg body weight in Glu-*CB*_*1*_-WT (Black circles, n=18) and Glu-*CB*_*1*_-KO (Open circles, n=11), GABA-*CB*_*1*_-WT (Black circles, n=6) and GABA-*CB*_*1*_-KO (Open circles, n=10), GFAP-*CB*_*1*_-WT (Black circles, n=7) and GFAP-*CB*_*1*_-KO (Open circles, n=11), D_1_-*CB*_*1*_-WT (Black circles, n=5) and D_1_-*CB*_*1*_-KO(Open circles, n=7). **i**, Cumulative water intake after 24-hour water deprivation in stop-*CB*_*1*_ (Open squares, n=8) and GABA-*CB*_*1*_-RS (Black squares, n=8). **j**, Cumulative water intake after IP NaCl, 10ml/kg body weight in stop-*CB*_*1*_ (Open squares, n=10) and GABA-*CB*_*1*_-RS (Black squares, n=8). All data are showed as mean ± s.e.m, and were statistically analyzed by the two-way repeated measurements ANOVA.

**Supplementary figure 3.**
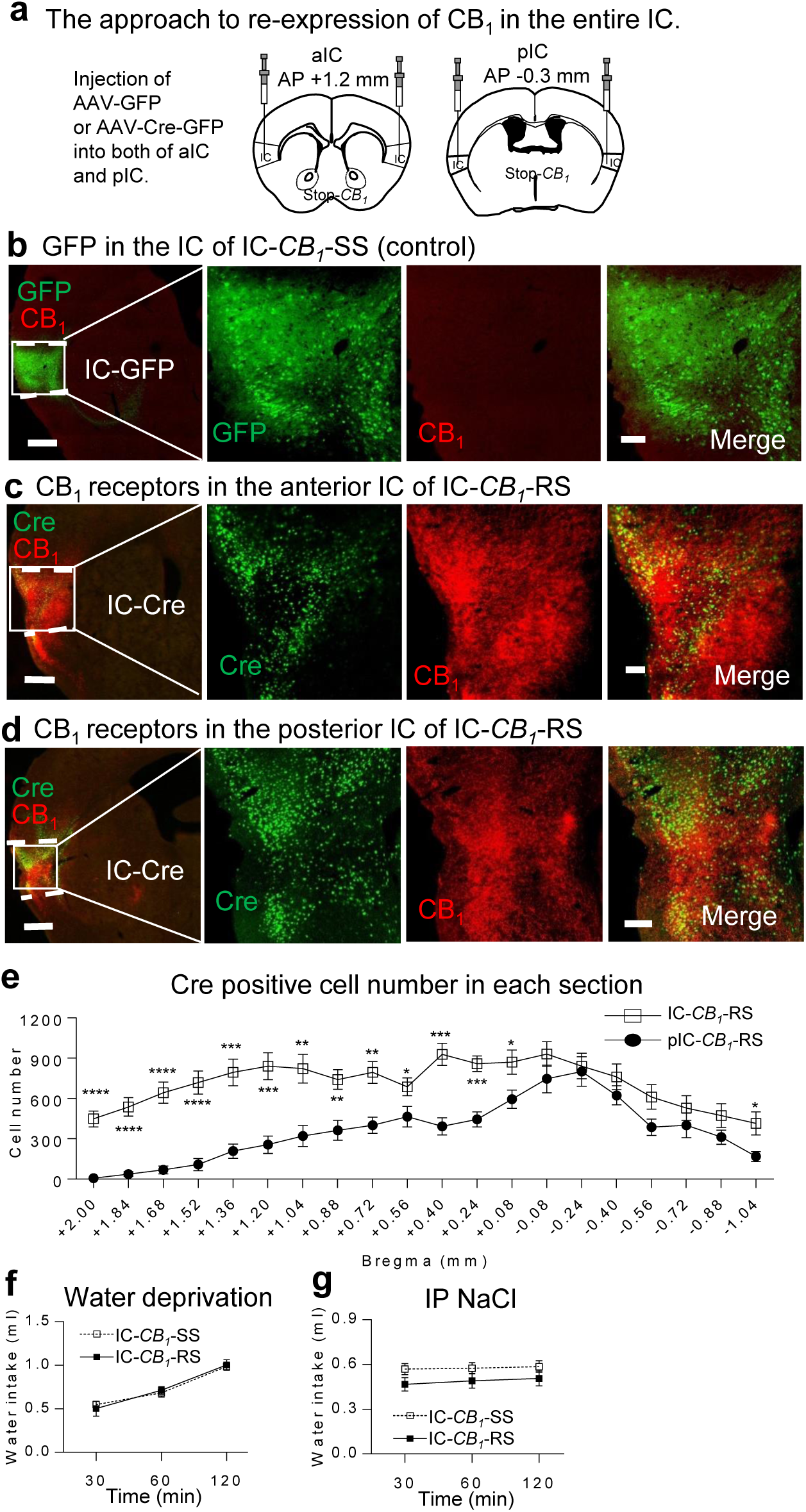
Re-expression of CB_1_ in the entire IC did not affect stimulated water intake. **a**, Schematic representation of CB_1_ rescue approach in the entire IC of Stop-*CB*_*1*_ mice. **b-d**, CB_1_ (red) immunostaining in the IC of IC-*CB*_*1*_-SS and IC-*CB*_*1*_-RS, respectively. Scale bar, 500μm and 100 (Amplified images) μm. **e**, Cre positive cell number in sequential brain sections of the IC-*CB*_*1*_-RS (Open squares, n=9) and pIC-*CB*_*1*_-RS (Black circles, n=8; the pIC data in the supplementary Figure 4). **f-g**, Cumulative water intake of IC-*CB*_*1*_-SS (Open squares) and IC-*CB*_*1*_-RS (Black squares) mice after 24-hour water deprivation (IC-*CB*_*1*_-SS n=8, IC-*CB*_*1*_-RS n=8) or IP 1M NaCl, 10ml/kg body weight (IC-*CB*_*1*_-SS n=9, IC-*CB*_*1*_-RS n=9). All data are showed as mean ± s.e.m. The data of Cre positive cell number were statistically analyzed by the two-tailed Student’s t-test. *\*P* < 0.05, *\*\*P* < 0.01, *\*\*\*P* < 0.001 *\*\*\*\**P < 0.0001.

**Supplementary figure 4.**
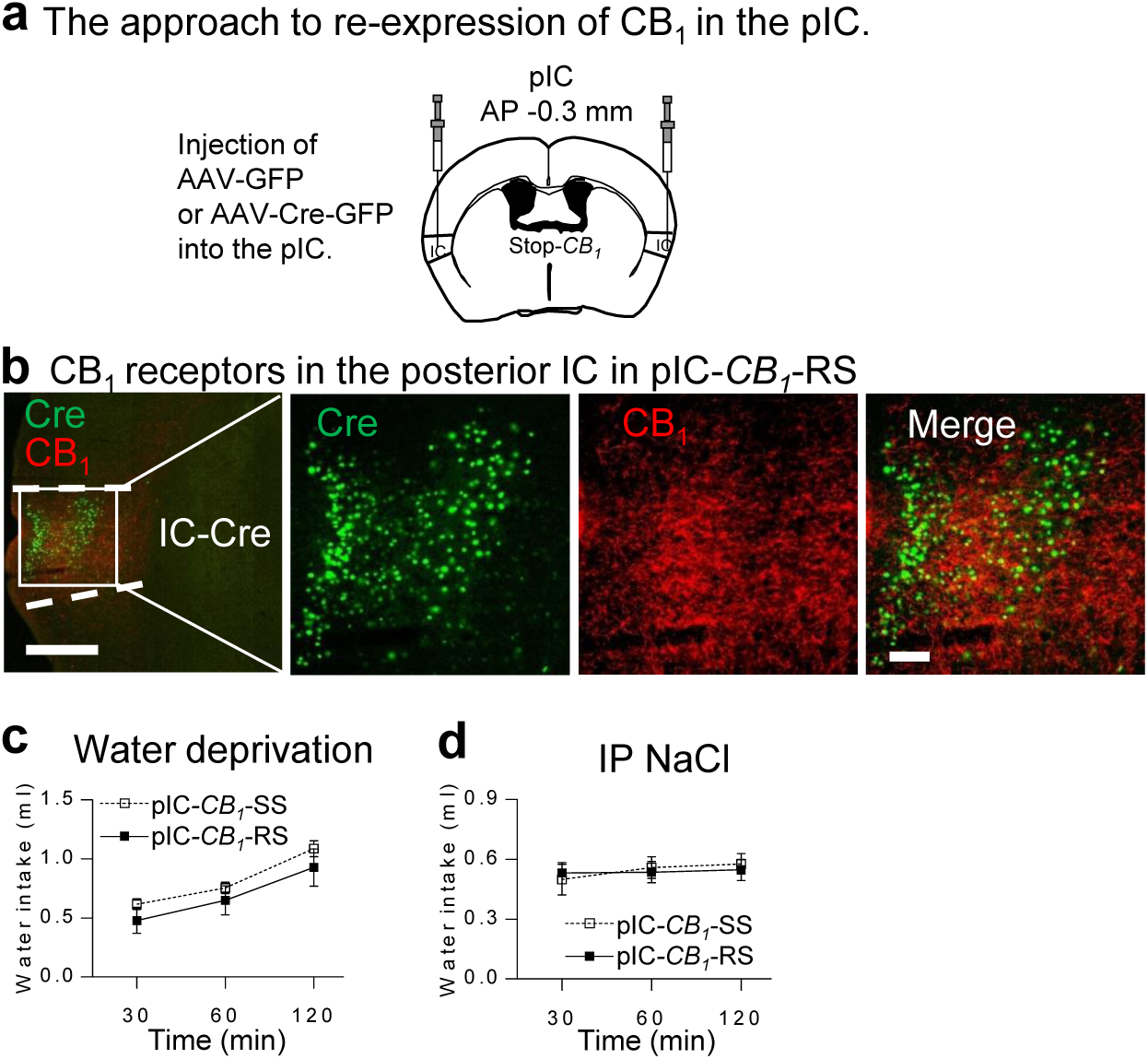
Re-expression of CB_1_ in the posterior IC did not affect stimulated water intake. **a**, Schematic representation of CB_1_ rescue approach in the posterior IC of Stop-*CB*_*1*_ mice. **b**, CB_1_ (red) immunostaining in the IC of pIC-*CB*_*1*_-RS. Scale bar, 500μm and 100 (Amplified images) μm. **c-d**, Cumulative water intake of pIC-*CB*_*1*_-SS (Open squares) and pIC-*CB*_*1*_-RS (Black squares) mice after 24-hour water deprivation (pIC-*CB*_*1*_-SS n=9, pIC-*CB*_*1*_-RS n=7) or IP 1M NaCl, 10ml/kg body weight (pIC-*CB*_*1*_-SS n=9, pIC-*CB*_*1*_-RS n=8). All data are showed as mean ± s.e.m, and were statistically analyzed by the two-way repeated measurements ANOVA.

**Supplementary figure 5.**
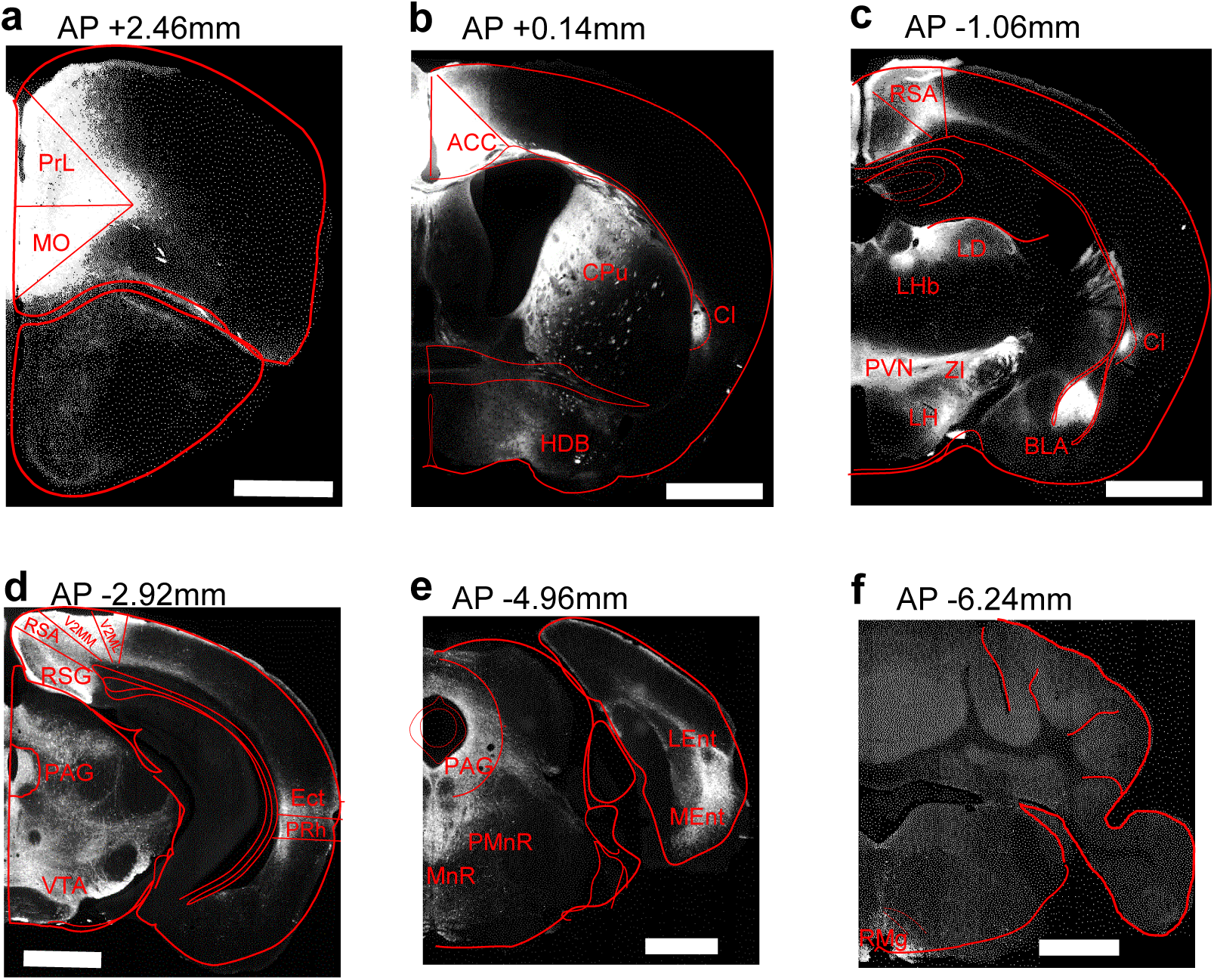
Brain-wide ACC neural projections revealed by injection of AAV-CaMKIIα-GFP into the ACC. **a**, Brain section at AP+2.46mm, PrL (Prelimbic cortex), MO (Medial obital cortex). **b**, Brain section at AP+0.14mm, CPu (Caudate putamen), CI (Claustrum), HDB (nucleus of the horizontal limb of the diagonal band). **c**, Brain section at AP-1.06mm, RSA (retrosplenial agranular cortex), LD (laterodorsal thalamic nucleus), LHb (Lateral habenula), PVN (paraventricular hypothalamic nucleus), ZI (zona incerta), LH (lateral hypothalamic area), BLA (basolateral amygdala). **d**, Brain section at AP-2.92mm, RSA (retrosplenial agranular cortex), RSG (retrosplenial granular cortex), V2MM (secondary visual cortex, mediomedial area), V2ML(secondary visual cortex, mediolateral area), PAG (periaqueductal gray), VTA (ventral tegmental area), Ect (ectorhinal cortex), PRh (perirhinal cortex). **e**, Brain section at AP-4.96mm, PAG (periaqueductal gray), MnR (median raphe nucleus), PMnR (paramedian raphe nucleus), LEnt (lateral entorhinal cortex), MEnt (medial entorhinal cortex). **f**, Brain section at AP-6.24mm, RMg (raphe magnus nucleus). Scale bar of a-f, 1 mm.

**Supplementary table 1.** Statistical details related to figures 1-3.

| Figure | Experiment, sample, size (n) | Analysis (post-hoc test) | Factors analyzed | F-ratios | P values |
| --- | --- | --- | --- | --- | --- |
| 1a | Water deprivation<br>*CB<sub>1</sub>*-WT (10) *CB<sub>1</sub>*-KO (8) | Two-way repeated measures ANOVA | Genotype and time | Interaction F (2, 32) = 2.254 | P = 0.1213 |
|  |  |  |  | Time F (2, 32) = 18.59 | P < 0.0001 |
|  |  |  |  | Genotype F (1, 16) = 19.49 | P = 0.0004 |
| 1b | IP 1M NaCl 10ml/kg<br>*CB<sub>1</sub>*-WT (10) *CB<sub>1</sub>*-KO (8) | Two-way repeated measures ANOVA | Genotype and time | Interaction F (2, 32) = 0.7755 | P = 0.4689 |
|  |  |  |  | Time F (2, 32) = 111.1 | P < 0.0001 |
|  |  |  |  | Genotype F (1, 16) = 16.02 | P = 0.0010 |
| 1c | ICV NaCl<br>*CB<sub>1</sub>*-WT (13) *CB<sub>1</sub>*-KO (10) | Two-way ANOVA | Genotype and dose | Interaction F (3, 84) = 2.776 | P = 0.0463 |
|  |  |  |  | Dose F (3, 84) = 19.55 | P < 0.0001 |
|  |  |  |  | Genotype F (1, 84) = 9.189 | P = 0.0032 |
| 1d | ICV ANG<br>*CB<sub>1</sub>*-WT (11) *CB<sub>1</sub>*-KO (13) | Two-way ANOVA | Genotype and dose | Interaction F (3, 88) = 3.292 | P = 0.0243 |
|  |  |  |  | Dose F (3, 88) = 79.76 | P < 0.0001 |
|  |  |  |  | Genotype F (1, 88) = 33.11 | P < 0.0001 |
| 1e | Water deprivation<br>C57BL/6 Vehicle (9)<br>C57BL/6 Rimonabant<br>3mg/kg (10) | Two-way repeated measures ANOVA | Drug and time | Interaction F (2, 34) = 6.461 | P = 0.0042 |
|  |  |  |  | Time F (2, 34) = 53.81 | P < 0.0001 |
|  |  |  |  | Drug F (1, 17) = 14.97 | P = 0.0012 |
| 1f | IP 1.5M NaCl 10ml/kg<br>C57BL/6 Vehicle (6)<br>C57BL/6 Rimonabant<br>3mg/kg (7) | Two-way repeated measures ANOVA | Drug and time | Interaction F (2, 22) = 0.9549 | P = 0.4002 |
|  |  |  |  | Time F (2, 22) = 27.89 | P < 0.0001 |
|  |  |  |  | Genotype F (1, 11) = 7.492 | P = 0.0193 |
| 1g | Water deprivation<br>Stop-*CB<sub>1</sub>*(9) *CB<sub>1</sub>*-RS (12) | Two-way repeated measures ANOVA | Genotype and time | Interaction F (2, 40) = 1.343 | P = 0.2726 |
|  |  |  |  | Time F (2, 40) = 80.70 | P < 0.0001 |
|  |  |  |  | Genotype F (1, 20) = 13.66 | P = 0.0014 |
| 1h | IP 1M NaCl 10ml/kg<br>Stop-*CB<sub>1</sub>*(10)<br>*CB<sub>1</sub>*-RS (11) | Two-way repeated measures ANOVA | Genotype and time | Interaction F (2, 38) = 0.1556 | P = 0.8564 |
|  |  |  |  | Time F (2, 38) = 14.88 | P < 0.0001 |
|  |  |  |  | Genotype F (1, 19) = 4.395 | P = 0.0497 |
| 1i | Water deprivation<br>Glu-*CB<sub>1</sub>*-SS (11)<br>Glu-*CB<sub>1</sub>*-RS (11) | Two-way repeated measures ANOVA | Genotype and time | Interaction F (2, 40) = 5.614 | P = 0.0071 |
|  |  |  |  | Time F (2, 40) = 103.9 | P < 0.0001 |
|  |  |  |  | Genotype F (1, 20) = 7.918 | P = 0.0107 |
| 1j | IP 1M NaCl 10ml/kg<br>Glu-*CB<sub>1</sub>*-SS (11)<br>Glu-*CB<sub>1</sub>*-RS (11) | Two-way repeated measures ANOVA | Genotype and time | Interaction F (2, 40) = 2.098 | P = 0.1360 |
|  |  |  |  | Time F (2, 40) = 14.26 | P < 0.0001 |
|  |  |  |  | Genotype F (1, 20) = 9.155 | P = 0.0067 |
| 1k | ICV NaCl<br>Glu-*CB<sub>1</sub>*-SS (13)<br>Glu-*CB<sub>1</sub>*-RS (11) | Two-way<br>ANOVA | Genotype<br>and dose | Interaction F (3, 84) = 3.132 | P = 0.0298 |
|  |  |  |  | Dose F (3, 84) = 8.909 | P < 0.0001 |
|  |  |  |  | Genotype F (1, 84) = 11.41 | P = 0.0011 |
| 1l | ICV ANG<br>Glu-*CB<sub>1</sub>*-SS (15)<br>Glu-*CB<sub>1</sub>*-RS (13) | Two-way<br>ANOVA | Genotype<br>and dose | Interaction F (3, 104) = 1.026 | P = 0.3845 |
|  |  |  |  | Dose F (3, 104) = 100.8 | P < 0.0001 |
|  |  |  |  | Genotype F (1, 104) = 6.506 | P = 0.0122 |
| 2d | Water deprivation<br>ACC-*CB<sub>1</sub>*-SS (17)<br>ACC-*CB<sub>1</sub>*-RS (20) | Two-way<br>repeated<br>mesures<br>ANOVA | Genotype<br>and time | Interaction F (2, 70) = 3.598 | P = 0.0325 |
|  |  |  |  | Time F (2, 70) = 182.5 | P < 0.0001 |
|  |  |  |  | Genotype F (1, 35) = 20.47 | P < 0.0001 |
| 2e | IP 1M NaCl 10ml/kg<br>ACC-*CB<sub>1</sub>*-SS (18)<br>ACC-*CB<sub>1</sub>*-RS (20) | Two-way<br>repeated<br>mesures<br>ANOVA | Genotype<br>and time | Interaction F (2, 72) = 0.1684 | P = 0.8454 |
|  |  |  |  | Time F (2, 72) = 26.92 | P < 0.0001 |
|  |  |  |  | Genotype F (1, 36) = 4.432 | P = 0.0423 |
| 3f | Water deprivation<br>ACC-BLA-*CB<sub>1</sub>*-SS (10)<br>ACC-BLA-*CB<sub>1</sub>*-RS (12) | Two-way<br>repeated<br>mesures<br>ANOVA | Genotype<br>and time | Interaction F (2, 40) = 0.2143 | P = 0.8080 |
|  |  |  |  | Time F (2, 40) = 59.32 | P < 0.0001 |
|  |  |  |  | Genotype F (1, 20) = 7.133 | P = 0.0147 |
| 3g | IP 1M NaCl 10ml/kg<br>ACC-BLA-*CB<sub>1</sub>*-SS (10)<br>ACC-BLA-*CB<sub>1</sub>*-RS (12) | Two-way<br>repeated<br>mesures<br>ANOVA | Genotype<br>and time | Interaction F (2, 40) = 1.556 | P = 0.2235 |
|  |  |  |  | Time F (2, 40) = 16.58 | P < 0.0001 |
|  |  |  |  | Genotype F (1, 20) = 10.28 | P = 0.0044 |

**Supplementary table 2.** Statistical details related to supplementary figures 1-4.

| Supplementary figure | Experiment, sample, size (n) | Analysis (post-hoc test) | Factors analyzed | F-ratios | P values |
| --- | --- | --- | --- | --- | --- |
| 1a | Daily water intake<br>*CB<sub>1</sub>*-WT (9)<br>*CB<sub>1</sub>*-KO (8) | Two-way ANOVA | Genotype and day | Interaction F (6, 105) = 0.2138 | P = 0.9717 |
|  |  |  |  | Days F (6, 105) = 0.3389 | P = 0.9149 |
|  |  |  |  | Genotype F (1, 105) = 2.150 | P = 0.1455 |
| 1b | Water intake in a day after water deprivation<br>*CB<sub>1</sub>*-WT (10)<br>*CB<sub>1</sub>*-KO (8) | Unpaired *t*-test |  |  | P = 0.0500 |
| 1c | Food intake in a day after water deprivation<br>*CB<sub>1</sub>*-WT (10)<br>*CB<sub>1</sub>*-KO (8) | Unpaired *t*-test |  |  | P = 0.5666 |
| 1d | IP NaCl dose response<br>*CB<sub>1</sub>*-WT (10)<br>*CB<sub>1</sub>*-KO (8) | Unpaired *t*-test |  |  | Saline P=0.1243 |
|  |  |  |  |  | 0.3M P=0.0130 |
|  |  |  |  |  | 0.5M P=0.1721 |
|  |  |  |  |  | 0.75M P=0.0226 |
|  |  |  |  |  | 1M P=0.0010 |
| 1e | Body water composition<br>*CB<sub>1</sub>*-WT (8)<br>*CB<sub>1</sub>*-KO (9) | Two-way ANOVA |  | Interaction F (1, 30) = 0.01329 | P = 0.9090 |
|  |  |  |  | Days F (1, 30) = 23.10 | P < 0.0001 |
|  |  |  |  | Genotype F (1, 30) = 1.242 | P = 0.2739 |
| 1f | Plasma osmolality<br>*CB<sub>1</sub>*-WT (8)<br>*CB<sub>1</sub>*-KO (8) | Unpaired *t*-test |  |  | P = 0.2496 |
| 2a | Water deprivation<br>Glu-*CB<sub>1</sub>*-WT (18)<br>Glu-*CB<sub>1</sub>*-KO (11) | Two-way repeated measures ANOVA | Genotype and time | Interaction F (2, 54) = 1.410 | P = 0.2529 |
|  |  |  |  | Time F (2, 54) = 109.5 | P < 0.0001 |
|  |  |  |  | Genotype F (1, 27) = 1.673 | P = 0.2068 |
| 2b | Water deprivation<br>GABA-*CB<sub>1</sub>*-WT (6)<br>GABA-*CB<sub>1</sub>*-KO (10) | Two-way repeated measures ANOVA | Genotype and time | Interaction F (2, 28) = 0.7316 | P = 0.4901 |
|  |  |  |  | Time F (2, 28) = 119.7 | P < 0.0001 |
|  |  |  |  | Genotype F (1, 14) = 2.026 | P = 0.1766 |
| 2c | Water deprivation<br>GFAP-*CB<sub>1</sub>*-WT (7)<br>GFAP-*CB<sub>1</sub>*-KO (11) | Two-way repeated measures ANOVA | Genotype and time | Interaction F (2, 32) = 0.8915 | P = 0.4200 |
|  |  |  |  | Time F (2, 32) = 46.29 | P < 0.0001 |
|  |  |  |  | Genotype F (1, 16) = 0.07548 | P = 0.7870 |
| 2d | Water deprivation<br>D1R-*CB<sub>1</sub>*-WT (5)<br>D1R-*CB<sub>1</sub>*-KO (7) | Two-way repeated measures ANOVA | Genotype and time | Interaction F (2, 20) = 0.6907 | P = 0.5128 |
|  |  |  |  | Time F (2, 20) = 9.770 | P = 0.0011 |
|  |  |  |  | Genotype F (1, 10) = 0.7693 | P = 0.4010 |
| 2e | IP 1M NaCl 10ml/kg<br>Glu-*CB<sub>1</sub>*-WT (18)<br>Glu-*CB<sub>1</sub>*-KO (11) | Two-way repeated measures ANOVA | Genotype and time | Interaction F (2, 54) = 0.4799 | P = 0.6215 |
|  |  |  |  | Time F (2, 54) = 13.36 | P < 0.0001 |
|  |  |  |  | Genotype F (1, 27) = 0.06145 | P = 0.8061 |
| 2f | IP 1M NaCl 10ml/kg<br>GABA-*CB<sub>1</sub>*-WT (6)<br>GABA-*CB<sub>1</sub>*-KO (10) | Two-way repeated measures ANOVA | Genotype and time | Interaction F (2, 28) = 0.6378 | P = 0.5360 |
|  |  |  |  | Time F (2, 28) = 10.47 | P = 0.0004 |
|  |  |  |  | Genotype F (1, 14) = 2.738 | P = 0.1202 |
| 2g | IP 1M NaCl 10ml/kg<br>GFAP-*CB<sub>1</sub>*-WT (7)<br>GFAP-*CB<sub>1</sub>*-KO (11) | Two-way repeated measures ANOVA | Genotype and time | Interaction F (2, 32) = 3.281 | P = 0.0506 |
|  |  |  |  | Time F (2, 32) = 22.50 | P < 0.0001 |
|  |  |  |  | Genotype F (1, 16) = 0.06804 | P = 0.7975 |
| 2h | IP 1M NaCl 10ml/kg<br>D1R-*CB<sub>1</sub>*-WT (5)<br>D1R-*CB<sub>1</sub>*-KO (7) | Two-way repeated measures ANOVA | Genotype and time | Interaction F (2, 20) = 0.1803 | P = 0.8364 |
|  |  |  |  | Time F (2, 20) = 7.256 | P = 0.0043 |
|  |  |  |  | Genotype F (1, 10) = 0.5358 | P = 0.4810 |
| 2i | Water deprivation<br>Stop-*CB<sub>1</sub>* (8)<br>GABA-*CB<sub>1</sub>*-RS (8) | Two-way repeated measures ANOVA | Genotype and time | Interaction F (2, 28) = 0.8169 | P = 0.4520 |
|  |  |  |  | Time F (2, 28) = 56.11 | P < 0.0001 |
|  |  |  |  | Genotype F (1, 14) = 0.2042 | P = 0.6583 |
| 2j | IP 1M NaCl 10ml/kg<br>Stop-*CB<sub>1</sub>* (10)<br>GABA-*CB<sub>1</sub>*-RS (8) | Two-way repeated measures ANOVA | Genotype and time | Interaction F (2, 32) = 1.099 | P = 0.3454 |
|  |  |  |  | Time F (2, 32) = 15.22 | P < 0.0001 |
|  |  |  |  | Genotype F (1, 16) = 2.626 | P = 0.1246 |
| 3e | Cre positive cell number<br>IC-*CB<sub>1</sub>*-RS (9)<br>pIC-*CB<sub>1</sub>*-RS (8) | Unpaired t-test | Bregema(mm) | +2.00 | P < 0.0001 |
|  |  |  |  | +1.84 | P < 0.0001 |
|  |  |  |  | +1.68 | P < 0.0001 |
|  |  |  |  | +1.52 | P < 0.0001 |
|  |  |  |  | +1.36 | P = 0.0002 |
|  |  |  |  | +1.20 | P = 0.0003 |
|  |  |  |  | +1.04 | P = 0.0021 |
|  |  |  |  | +0.88 | P = 0.0024 |
|  |  |  |  | +0.72 | P = 0.0016 |
|  |  |  |  | +0.56 | P = 0.0425 |
|  |  |  |  | +0.40 | P = 0.0001 |
|  |  |  |  | +0.24 | P = 0.0001 |
|  |  |  |  | +0.08 | P = 0.0296 |
|  |  |  |  | -0.08 | P = 0.1974 |
|  |  |  |  | -0.24 | P = 0.7861 |
|  |  |  |  | -0.40 | P = 0.2704 |
|  |  |  |  | -0.56 | P = 0.0629 |
|  |  |  |  | -0.72 | P = 0.3328 |
|  |  |  |  | -0.88 | P = 0.1490 |
|  |  |  |  | -1.04 | P = 0.0229 |
| 3f | Water deprivation<br>IC-CB<sub>1</sub>-SS (8)<br>IC-CB<sub>1</sub>-RS (8) | Two-way<br>repeated<br>measures<br>ANOVA | Genotype and<br>time | Interaction F (2, 28) = 0.5276 | P = 0.5958 |
|  |  |  |  | Time F (2, 28) = 73.13 | P < 0.0001 |
|  |  |  |  | Genotype F (1, 14) =<br>4.499e-015 | P > 0.9999 |
| 3g | IP 1M NaCl 10ml/kg<br>IC-CB<sub>1</sub>-SS (9)<br>IC-CB<sub>1</sub>-RS (9) | Two-way<br>repeated<br>measures<br>ANOVA | Genotype and<br>time | Interaction F (2, 32) = 2.137 | P = 0.1346 |
|  |  |  |  | Time F (2, 32) = 13.32 | P < 0.0001 |
|  |  |  |  | Genotype F (1, 16) = 2.095 | P = 0.1671 |
| 4c | Water deprivation<br>pIC-CB<sub>1</sub>-SS (9)<br>pIC-CB<sub>1</sub>-RS (7) | Two-way<br>repeated<br>measures<br>ANOVA | Genotype and<br>time | Interaction F (2, 28) = 0.2249 | P = 0.8000 |
|  |  |  |  | Time F (2, 28) = 70.61 | P < 0.0001 |
|  |  |  |  | Genotype F (1, 14) = 1.216 | P = 0.2888 |
| 4d | IP 1M NaCl 10ml/kg<br>pIC-CB<sub>1</sub>-SS (9)<br>pIC-CB<sub>1</sub>-RS (8) | Two-way<br>repeated<br>measures<br>ANOVA | Genotype and<br>time | Interaction F (2, 45) = 0.1708 | P = 0.8435 |
|  |  |  |  | Time F (2, 45) = 0.3459 | P = 0.7095 |
|  |  |  |  | Genotype F (1, 45) = 0.02060 | P = 0.8865 |

**Supplementary Video 1**. Whole-brain mapping of ACC neural projections by injection of AAV-CaMKIIα-GFP into ACC.

**Supplementary Video 2**. Whole-brain mapping of CB_1_ receptors’ distribution in ACC-*CB*_*1*_-RS mice.

